# Model-based integration of genomics and metabolomics reveals SNP functionality in *Mycobacterium tuberculosis*

**DOI:** 10.1101/555763

**Authors:** Ove Øyås, Sonia Borrell, Andrej Trauner, Michael Zimmermann, Julia Feldmann, Sebastien Gagneux, Jörg Stelling, Uwe Sauer, Mattia Zampieri

## Abstract

Human tuberculosis is caused by members of the *Mycobacterium tuberculosis* complex (MTBC) and presents variable disease outcomes. The variation has primarily been attributed to host and environmental factors, but recent evidence indicates an additional role of genetic diversity among MTBC clinical strains. Here, we used metabolomics to unravel the potential role of genetic variations in conferring strain-specific adaptive capacity and vulnerability. To systematically identify functionality of single nucleotide polymorphisms (SNPs), we developed a constraint-based approach that integrates metabolomic and genomic data. Model-based predictions were systematically tested against independent metabolome data; they correctly classified SNP effects in pyruvate kinase and suggested a genetic basis for strain-specific sensitivity to the antibiotic *para*-aminosalicylic acid. Our method is broadly applicable to mutations in enzyme-encoding genes across microbial life, opening new possibilities for identifying strain-specific metabolic vulnerabilities that could lead to more selective treatment strategies.

---

The *Mycobacterium tuberculosis* complex (MTBC) consists of closely related bacteria that are etiological agents of tuberculosis (TB) in humans and animals. Systematic genotyping and whole-genome sequencing resolved the landscape of evolved genetic differences in MTBC and distinguished seven phylogenetic lineages adapted to humans that feature limited genetic diversity^1,2,3,4^. Given the low mutation rate, strictly clonal reproduction, and the limited genetic diversity of the MTBC^5,6^, the variable outcomes of TB infection and disease as well as antibiotic treatments have been mainly attributed to host diversity rather than bacterial variations^7^. More recently, a growing body of evidence related genetic diversity among clinical MTBC strains to distinct geographic distributions^8^ and key selective advantages such as evolvability, transmissibility, and antibiotic tolerance^9,10^, suggesting that strain diversity may be an important factor^11,12^.

A key element of MTBC’s pathogenic success is its ability to evade the host immune response^13^ and to survive long periods of hypoxia, nutrient limitation, and oxidative or nitro-oxidative stresses^14,15,16^, which requires evolved plasticity for different metabolic responses. It is an open question how much of MTBC’s genetic diversity confers strain-specific metabolic characteristics, ultimately affecting transmissibility, disease progression, and antibiotic tolerance. Indeed, evolution of metabolism has been demonstrated to play a key role for successful adaptation to complex and dynamic *in vivo* environments^17,18^, for bacterial drug tolerance^19,20,21^, and for survival during infection of macrophages^22,23,24,25^. Beyond analyses of individual laboratory strains, however, no systematic characterization and comparative analysis of intrinsic metabolic differences across human-adapted MTBC strains has been performed.

If the metabolic and other phenotypic diversity between MTBC strains contributes to and modulates pathogenicity, the next question is: which elements of the limited genetic diversity in the MTBC are responsible for phenotypic strain diversification? In many species, associations between observable traits and single-nucleotide polymorphisms (SNPs) have been detected through genome-wide association studies (GWASs)^26,27^. GWASs typically estimate correlations between individual SNPs and phenotypes such as fitness or resistance, but large sample sizes are necessary to achieve adequate statistical power and to identify genes responsible for complex phenotypes. This is particularly challenging for the MTBC, given the sporadic nature of genetic differences and the practical difficulties in systematically characterizing strain-specific phenotypes. Alternative computational solutions to establish causal links between genotype and phenotype, such as homology-based methods like SIFT^28^ or VIPUR^29^, often exhibit low specificity (i.e., high risk of false positives)^30^, and do not consider correlations between genetic variants. Moreover, we do not know of any validation of these methods in microbes. As a result, only a small proportion of SNPs in the MTBC have been mechanistically linked to phenotypes such as virulence or intrinsic drug resistance^31,32^.

To overcome the limitations of GWASs and homology-based predictions, we developed a computational approach that integrates strain-specific exometabolomes, genomes, and genome-scale metabolic networks into a single model. Application to 18 representative human-adapted MTBC strains allowed us to predict the effects of SNPs in enzyme-encoding genes on metabolic phenotypes and identify functional SNPs that associate with strain-specific metabolic vulnerabilities and resistance to antibiotics.

## Results

To investigate metabolic diversity in the human-adapted MTBC, we selected three strains each from six of the seven known MTBC lineages^8^, covering much of the intra-lineage diversity^33^ (**Fig. 1a**). Lineage 7 was only recently characterized^34^ and not included here because strains were not readily available. All 18 MTBC strains were grown in batch cultures with a modified 7H9 medium containing pyruvate as the main carbon source and without common additives such as Tween 80, oleic acid, and glycerol. This medium supports growth of all strains while simplifying the composition to facilitate downstream analysis. We collected supernatant samples from exponentially growing cultures over a period of five days and monitored cell growth by optical density. The supernatants were analyzed with a non-targeted metabolomics method using a time-of-flight mass spectrometer^35^.

**Figure 1:**
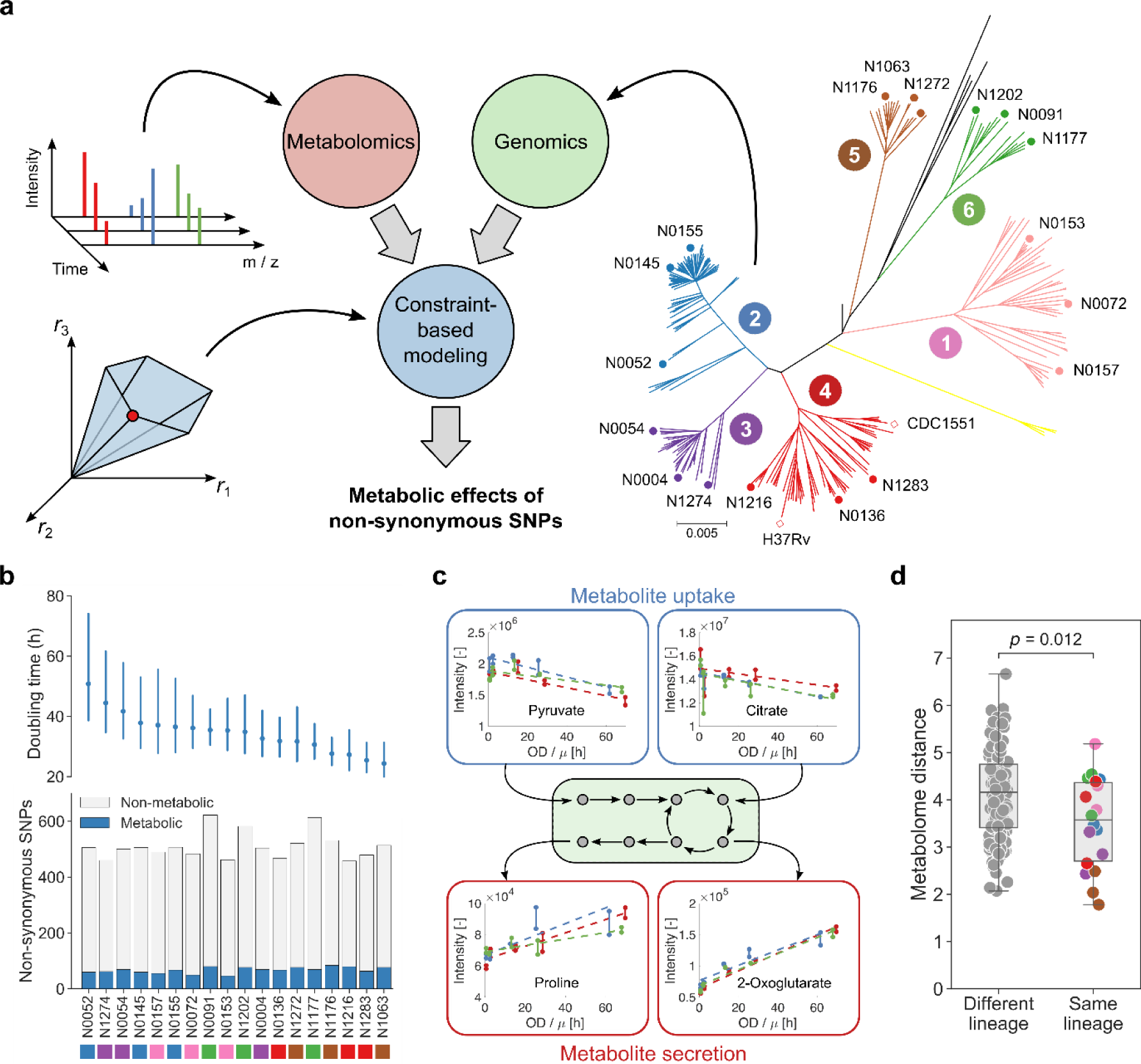
Connecting genetic and phenotypic diversity in MTBC. **a**, We integrated the genomes and dynamic exometabolomes of 18 human-adapted MTBC strains (three strains each from six lineages, as indicated in the phylogenetic tree) into a single constraint-based model that predicts the effects of non-synonymous SNPs on metabolic fluxes. **b**, Doubling times (circles) with 95% confidence intervals (lines) estimated from three pooled replicate experiments for each strain (above) and number of metabolic and non-metabolic non-synonymous SNPs (below) for all strains (colored by lineage). **c**, Relative metabolite uptake and secretion rates were inferred by fitting linear models (dashed lines) to dynamic exometabolomes. In this example, uptake of pyruvate and citrate and secretion of proline and 2-oxoglutarate are shown for strain N1202. Biological replicates (three for each strain) are indicated by color and technical replicates (two for each time point) are connected by solid lines. **d**, Metabolome distance (Euclidean distance between relative uptake and secretion rates inferred from exometabolomes) for all pairs of strains. Strains from the same lineage had more similar exometabolomes than strains from different lineages (*t*-test, *p* = 0.012, *n* = 153). Pairs of strains from the same lineage are colored according to the phylogenetic tree of panel a.

There are 4,717 non-synonymous SNPs in our 18 MTBC strains^36^, 566 of which could be mapped to enzyme-encoding genes in a genome-scale metabolic model^37^ (**Supplementary Tables 1 and 2**). Despite few SNPs in metabolic genes, the doubling times of our MTBC strains ranged from 25 to 50 hours (**Fig. 1b**), suggesting different adaptive metabolic strategies for the uptake and utilization of available nutrients. To link growth and metabolic phenotypes, we used the dynamic exometabolome profiles of 272 putative metabolites to derive relative nutrient uptake and byproduct secretion rates (**Fig. 1c and Supplementary Table 3**). Consistent with variation in doubling times, about 50% of the metabolites were differentially secreted or consumed in at least one of the 18 strains (maximum log_2_ fold change between strains larger than 2). Overall, the relative uptake and secretion rates of metabolites exhibited variation across strains by up to two orders of magnitude (**Supplementary Fig. 1**). The most significant (ANOVA, *q* ≤ 0.01) exometabolome differences across MTBC strains pertained to amino acids and intermediates of glycolysis and the tricarboxylic acid (TCA) cycle, suggesting divergent evolutionary strategies in central carbon and nitrogen metabolism (**Supplementary Tables 4 and 5**). Since the doubling time did not significantly (*q* ≤ 0.01) correlate with any of the metabolite uptake and secretion patterns (**Supplementary Fig. 1**), the metabolic diversity was unlikely due to indirect, growth-related effects but rather caused by genetic differences. Consistent with this hypothesis, we found that strains from the same lineage had significantly (*t*-test, *p* = 0.012, *n* = 153) more similar exometabolome profiles than strains from different lineages (**Fig. 1d**).

To identify genetic variations responsible for the observed metabolic differences among MTBC strains, we built on flux balance analysis (FBA)^38^, a constraint-based approach that predicts metabolic fluxes at steady state in a stoichiometric model of metabolic reactions by optimizing a cellular objective function. Here, we developed an FBA-based differential analysis to predict the effects of strain-specific SNPs via perturbation of the associated enzymes and propagation of this perturbation through the metabolic network (**Fig. 2a**). Specifically, we considered only non-synonymous (missense) SNPs in enzyme-encoding genes that potentially reduced enzymatic activity. We represented all 18 MTBC strains by a single constraint-based model (**Fig. 2b**) that contained genome-scale metabolic networks specific for each strain (incorporating gene deletions and experimentally determined bounds on growth and carbon source uptake rates), a virtual reference strain that allowed us to infer strain-specific differences, and strain-specific bounds on uptake and secretion (exchange) fluxes derived from exometabolome data. The model couples strains to each other through mass balances, SNP effects (a SNP must have the same effect in all affected strains), and measured ratios of exchange fluxes. A strain’s steady-state flux distribution, **r**_*i*_, (**Fig. 2c**) is then given by the reference strain flux distribution, **r**_ref_, and deviations from this reference that are either caused by SNP effects, **e**, and propagated through the network by a strain-specific structural sensitivity matrix^39^, **S**_*i*_, or due to effects that cannot be explained by metabolic SNPs, **u**_*i*_. To infer the unknowns **r**_*i*_, **r**_ref_, **e**, and **u**_*i*_, we sequentially minimized the sum of absolute values (*L*^1^ norm) of (i) unexplained flux differences between strains (**u**_*i*_), (ii) fluxes in the reference strain (**r**_ref_), and (iii) the SNP effects (**e**). This approach aimed to explain as much of the observed metabolic variation as possible by as few SNPs as possible while obtaining biologically reasonable flux distributions across the strains.

**Figure 2:**
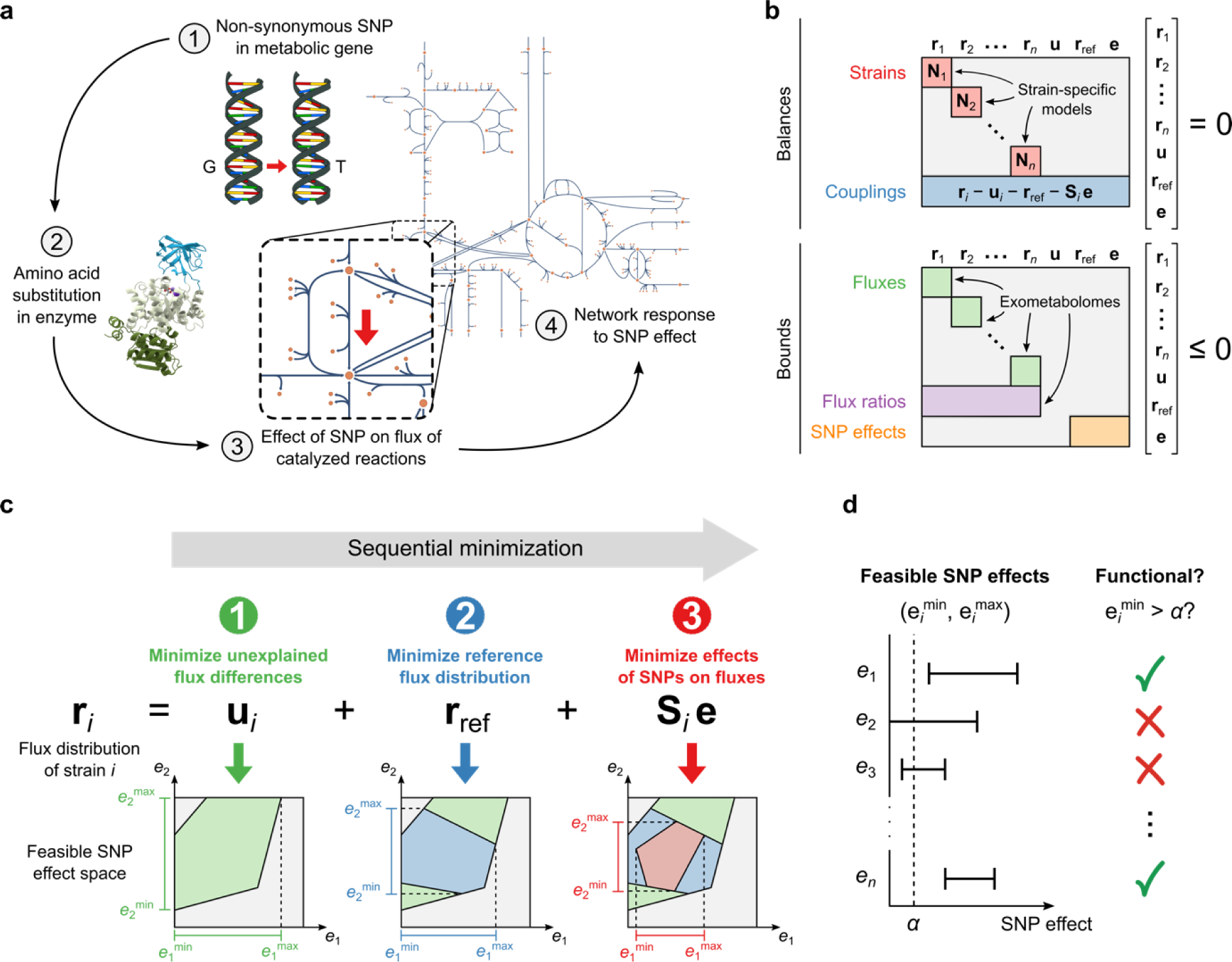
Predicting the metabolic effects of non-synonymous SNPs. **a**, Our constraint-based model integrates the metabolic effects of all non-synonymous SNPs in enzyme-encoding genes. Each SNP effect is represented in the following way: (1) a non-synonymous SNP causes (2) an amino acid substitution in an enzyme, possibly leading to reduced enzyme activity and (3) reduced flux through catalyzed reactions, and (4) the metabolic network responds by adjusting fluxes to a new steady state. The last two steps, as defined by structural sensitivity analysis^39^, are represented in the model. **b**, Our constraint-based model consists of balances and bounds, which are equality and inequality constraints, respectively. The constrained variables are the strain-specific flux distributions (**r**_1_, **r**_2_,…, **r**_*n*_), fluxes in a reference strain shared by all strains (**r**_ref_), unexplained flux differences between strains (**u**), and flux differences between strains explained by SNP effects (**e**). The balance constraints are the steady-state mass balances of strain-specific models (red) and couplings that ensure that differences between strain-specific fluxes and reference fluxes are accounted for by unexplained effects and SNP effects (blue). Bounds are applied to all fluxes in the strain-specific models, including bounds on exchange fluxes obtained from exometabolomes (green), the ratio of exchange fluxes between pairs of strains, also from exometabolomes (purple), and the size of SNP effects relative to their affected reference fluxes (orange). **c**, SNP effects are predicted by sequentially minimizing (1) unexplained flux differences between strains, (2) reference fluxes, and (3) flux differences explained by SNPs. The minimum obtained after each step is used to constrain the model, reducing the feasible space of solutions before proceeding to the next step. In this example, the feasible space of two SNP effects (*e*_1_ and *e*_2_) is shown after each minimization with feasible SNP effect ranges indicated along the axes. **d**, Feasible ranges of SNP effects (*e*_1_, *e*_2_,…, *e*_n_) can be compared directly and a chosen threshold, *α*, can be used to classify SNPs as functional or non-functional after each sequential minimization step. SNPs with a minimal effect larger than *α* are classified as functional.

A common problem of constraint-based models is that predicted flux distributions, or in our case, combinations of SNP effects, are not unique^40^. We therefore used flux variability analysis (FVA)^40^ to find the full range of possible values for all SNP effects across all possible optimal solutions. We classified a SNP as functional if its smallest possible effect in all optimal solutions was above a flux threshold *α*, and non-functional otherwise. (**Fig. 2d**). To avoid false positives, we chose the threshold value *α* = 10^-4^ mmol gDW^-1^ h^-1^ by systematically testing all possible values and selecting the one at which the number of SNPs classified as functional started decreasing more slowly with increasing (stricter) *α* (**Supplementary Fig. 2**). Results were robust to small changes in the threshold value (**Supplementary Figs. 2 and 3**).

**Figure 3:**
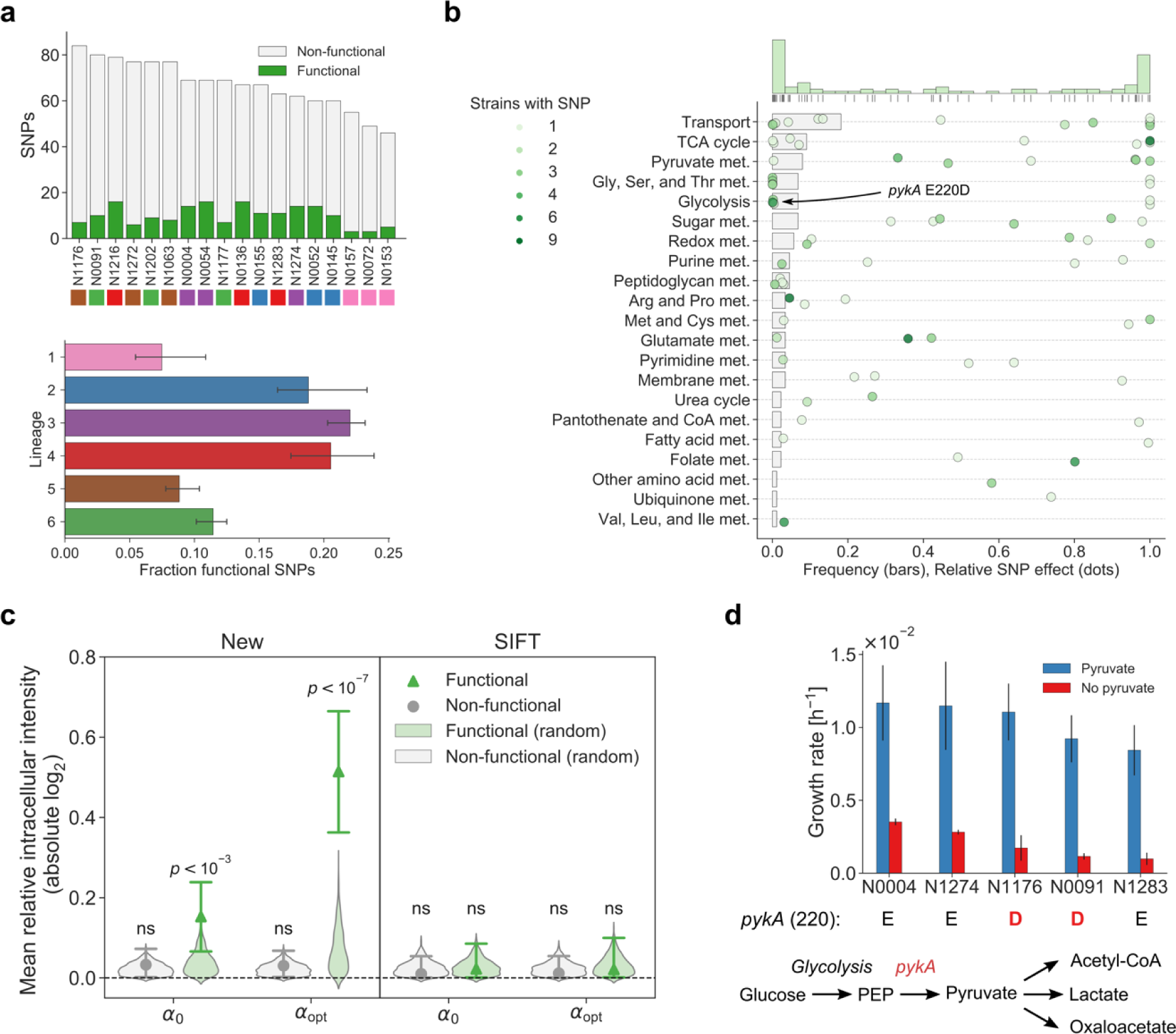
Predicted functional SNPs in the MTBC and validation of their metabolic effects. **a**, Number of non-synonymous SNPs classified as functional and non-functional in our 18 MTBC strains (above) and fraction of SNPs classified as functional by lineage (below). Lineages are indicated by color. Error bars indicate 95% confidence intervals from bootstrapping. **b**, Metabolic pathway distribution of functional SNPs and their relative effects on affected enzymes. Grey bars indicate the frequency of functional SNPs in each pathway. Each dot represents a functional SNP and indicates its minimal effect divided by the mean predicted reference flux of reactions catalyzed by affected enzymes (darker color indicates more strains bearing the mutations). **c**, Relative intracellular levels of metabolites participating in reactions catalyzed by enzymes affected by SNPs (ratios of affected metabolite levels between strains with and without SNPs). Mean absolute values are shown with 95% confidence intervals from bootstrapping and distributions of mean levels obtained from assigning SNPs randomly to strains while preserving the number of strains affected by each SNP. Results are shown for our new method and SIFT, for the threshold used for SNP classification (*α*) as well as a stricter threshold that optimizes the difference between functional and non-functional SNPs (*α*_opt_). The mean relative levels of functional SNPs predicted by our new method were significantly different from random distributions for *α* (*Z*-test, *p* = 2.0 × 10^−4^, *n* = 1,000) as well as for *α*_opt_(*Z*-test, *p* = 9.4 × 10^−8^, *n* = 1,000), but we did not find any significant difference using predictions from SIFT. **d**, Estimated growth rates of five strains grown in glucose medium with (blue) and without (red) supplemented pyruvate. Strains affected by the *pykA* E220D SNP are indicated below. One experiment was performed for each strain and condition. Error bars indicate two standard errors of the growth rate estimate. The pathway diagram shows *pykA* in its metabolic context.

Of the 566 non-synonymous SNPs in enzyme-encoding genes represented in the model, 88 (16%) were classified as functional. Notably, only 29 SNPs were predicted to be functional both by our constraint-based approach and SIFT (p>0.5, hypergeometric test) (**Supplementary Table 2**). Overall, strains from lineages 2, 3, and 4 had a slightly larger fraction of functional SNPs than strains from lineages 1, 5, and 6 (**Fig. 3a**). The functional SNPs affected 67 unique enzymes distributed across most metabolic pathways, and SNPs were most often predicted to either abolish the flux of a reaction (i.e., complete loss-of-function) or have a relatively minor effect under the tested condition (**Fig. 3b and Supplementary Table 2**). Metabolic pathways with many functional SNPs and large relative SNP effects included the TCA cycle, pyruvate metabolism, and glycolysis as well as glycine, serine, and threonine metabolism, consistent with our previous observation of significant differences in secretion rates of central metabolism intermediates and amino acids.

To systematically benchmark our predictions, we tested whether the levels of intracellular metabolites directly linked to enzymes with predicted functional SNPs exhibited larger variations across strains, than metabolites proximal to enzymes with non-functional SNPs. Our premise is that functional mutations affect enzyme kinetics (e.g. K_m_, K_cat_), resulting in local adaptive changes of metabolite levels^41^. We measured the intracellular levels of 294 putative metabolites in the 18 MTBC strains. For each SNP affecting an enzyme, we calculated the ratios of directly affected metabolite levels between strains with and without the SNP (**Supplementary Table 6**). On average, these ratios were significantly different between functional and non-functional SNPs (*t*-test, *p* = 2.8 × 10^−4^, *n* = 1,208). Moreover, the effects of functional SNPs were significantly different from distributions obtained by randomly assigning SNPs to strains (*Z*-test, *p* = 2.0 × 10^−4^, *n* = 1,000), whereas the ratios of non-functional SNPs were not (*Z*-test, *p* = 0.91, *n* = 1,000). This separation could be optimized by making the functional SNP classification more stringent, indicating that SNPs with larger predicted effects had a greater impact on intracellular metabolite levels (**Fig. 3c**). In contrast, functional SNPs predicted by SIFT^28^ did not achieve a significant separation for any classification threshold (**Supplementary Fig. 3)**. Thus, the analysis of intracellular metabolome data across the 18 MTBC strains independently supported our model-based classification of SNP functionality over homology-based predictions.

To validate our SNP functionality predictions more directly, we focused on the known functional E200D substitution in the glycolytic enzyme pyruvate kinase (PykA)^42,43^, a mutation that occurs in all animal-adapted MTBC strains and the human-adapted lineages 5 and 6, also known as *M. africanum*^44^. This SNP was shown to abolish the activity of PykA in animal-adapted strains, rendering them unable to grow on glucose as the sole carbon source^42,43^. Deleting *pykA* in the laboratory strain H37Rv also leads to glucose toxicity, presumably due to accumulation of methylglyoxal^45^. While our analysis classified the *pykA* E220D SNP as functional, it predicted only a relatively small flux effect under the tested conditions in E200D-carrying MTBC strains (**Fig. 3b**). To test this prediction, we grew five MTBC strains on 7H9 glucose medium with and without pyruvate. Clinical isolates with the E200D substitution were able to grow on glucose both with and without pyruvate, albeit to a lesser extent than strains without the SNP, in agreement with our prediction (**Fig. 3d**). Consistently, the intracellular levels of methylglyoxal were comparable in strains with and without the E200D SNP (**Supplementary Fig. 4**). Thus, our physiological data validate the model prediction of *pykA* E200D functionality, and indicate that under the tested conditions clinical isolates behave differently from the animal-adapted and laboratory strains.

**Figure 4:**
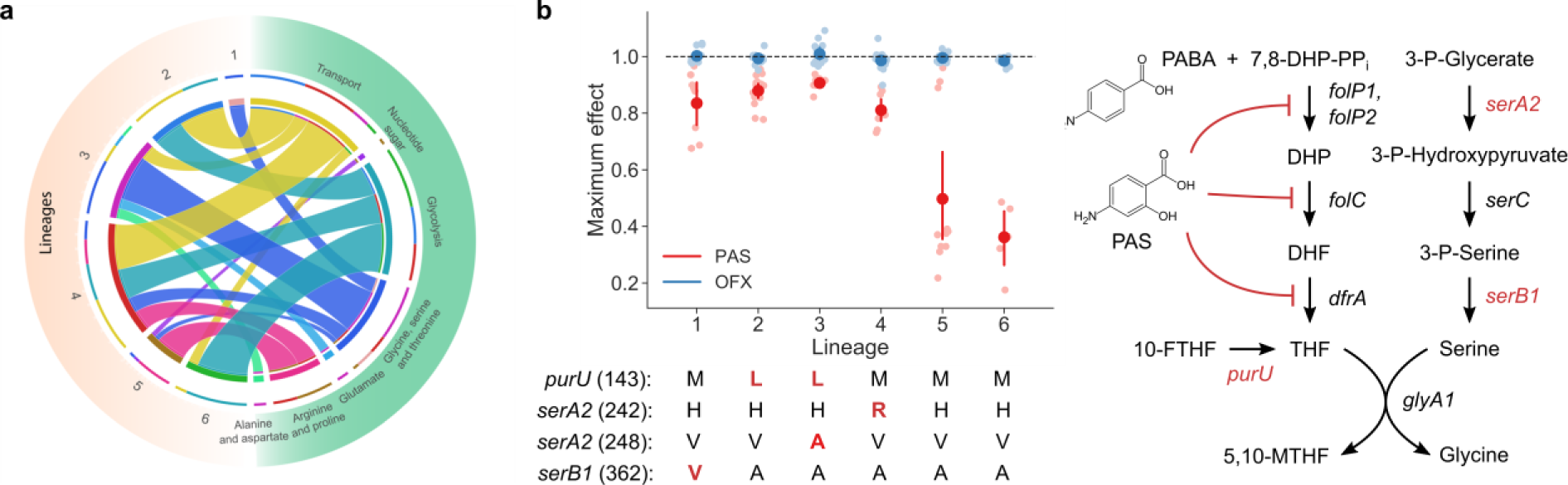
Predicted functional SNPs suggest causes of lineage-specific metabolic vulnerabilities in MTBC. **a**, Circos plot^65^ showing predicted synthetic lethal interactions involving enzymes affected by functional SNPs. Each ribbon connects an MTBC lineage to a metabolic subsystem and its size indicates the number of functional SNPs in the lineage that epistatically interact with an enzyme in the metabolic pathway. **b**, Sensitivity of strains to the antibiotics *para*-aminosalicylic acid (PAS) and ofloxacin (OFX). The plot shows the mean relative viability (viability at highest drug concentration relative to lowest) for each lineage with 95% confidence intervals from bootstrapping (dose response curves with number of replicates shown in **Supplementary Figs. 5 and 6**). Lineages 1-4 were significantly less sensitive to PAS than lineages 5 and 6 (*t*-test, *p* = 4.5 × 10^−13^, *n* = 55) while all strains were sensitive to OFX. Because of PAS-insensitive strains N1176, N1272, N009 and N1177, maximum drug effect was estimated rather than potency (IC_50_)^66^. The pathway diagram shows the mode of action of PAS and its connection to the enzymes affected by four functional SNPs in folate metabolism that could explain the observed differences in sensitivity.

Next, we systematically investigated whether the predicted functional SNPs could reveal strain-specific metabolic vulnerabilities. We used FBA to find all synthetic lethal pairs of gene deletions in which one of the genes contained at least one of our predicted functional SNPs (modeled as deletions in this analysis). The predicted epistatic interactions of SNP-affected enzymes differed between strains and were mostly with enzymes involved in glycolysis, amino acid biosynthesis, and nutrient transport (**Fig. 4a**). For example, the majority of the predicted epistatic interactions in lineages 1 and 3 involved enzymes in glycine, serine, and threonine metabolism, while strains from lineages 2, 4, and 6 were predicted to be more vulnerable to disruptions in glycolysis. Thus, our analysis suggests that MTBC genetic diversity can confer strain-specific sensitivity to the inhibition of metabolic functions.

Several functional SNPs conferring strain-specific vulnerabilities were found in pathways dependent on folate cofactors, such as glycine, and serine biosynthesis, also including purine, methionine and pantothenate metabolism (**Fig. 3b**). Since folate biosynthesis is a validated metabolic target for antibiotics^46^, we tested the functionality of SNPs by differential susceptibility of MTBC strains to inhibitors of folate metabolism. To this end, we determined the sensitivity of 15 of the 18 MTBC strains to *para*-aminosalicylic acid (PAS), an inhibitor of tetrahydrofolate (THF) biosynthesis and one of the antimycobacterial drugs currently used for multi-drug-resistant TB^47,48^. Although no strain carried the genetic determinants of PAS resistance^49^, strains from lineages 1-4 were significantly more susceptible to PAS than strains from lineages 5 and 6 (*t*-test, *p* = 4.5 × 10^−13^, *n* = 55) (**Fig. 4b**). We identified four predicted functional SNPs in three enzymes proximal to THF that could help explain these differences (**Fig. 4b**). These SNPs are present in sensitive strains; they affect *serA2* and *serB1* (needed for serine biosynthesis) and *purU* (which converts 10-formyl-THF to THF). As a control, all strains were equally sensitive to the antibiotic ofloxacin (OFX) (**Fig. 4b**). We speculate that, similar to previous observations in *Escherichia coli*^50^, recycling of 10-formyl-THF and 5,10-methylene-THF via *purU*, *serA2*, and *serB1* plays a key role in replenishing the THF pool to compensate for the effect of the antibiotic, and that the SNPs impair this role.

## Discussion

MTBC strains cluster into seven distinct phylogeographic lineages^2^ that have been proposed to be adapted to different host populations^51^. Current GWAS methods for relating genetic variation to phenotype, were successful in identifying clinically prevalent mutations that confer strong phenotypes such as drug resistance^31^. However, because the genetic diversity in MTBC strains is small compared to other pathogens^52,6^, such standard methods struggle to relate genetic variation to host adaptation. Moreover, it is unclear what the relevant phenotypes for host adaptation are. Because metabolic adaptability is crucial for MTBC strains to scavenge nutrients in harsh host environments^53^, we hypothesized that metabolic phenotypes could discriminate between MTBC strains and their clinical phenotypes. Indeed, despite their limited genetic diversity^51,6^, our data for 18 MTBC strains from six lineages demonstrate substantial lineage-specific phenotypic differences between MTBC strains, reflected in different growth rates and preferences for nutrient uptake and byproduct secretion, potentially revealing divergent metabolic specialization in MTBC lineages^54,55^. This suggests that integrating genome and metabolome profiling can help find strain-specific factors for host adaptation as well as metabolic vulnerabilities that are strain-specific.

To link metabolic phenotypes to genomic information, we developed a novel constraint-based method to identify functional SNPs affecting enzymes, that is, non-synonymous SNPs that are required to explain the experimental observations such as growth, uptake and secretion rates. The central concept of the method is to represent all strains and experimental information in a single constraint-based model to consider all possible dependencies between SNPs and their effects on metabolic phenotypes. This systematic data integration allowed us to predict the functionality of enzyme SNPs despite a relatively small sample size and restricted genetic variability. This computational strategy is complementary to large-scale GWAS and is broadly applicable to other organisms, in particular because of the large and rapidly increasing number of microbes for which sequencing information and genome-scale metabolic models are available. The current approach is restricted to enzymatic SNPs, but extensions could include control sequences of metabolic genes (e.g., by representing SNP-modulated enzyme concentrations) as well as other forms of genetic variability. Even with this limitation, our model-based prediction of functional SNPs has a key advantage over homology-based methods such as SIFT^28^: it investigates SNP functionality in the context of potential compensatory mechanisms (i.e., alternative metabolic pathways) and other genetic variations. This was crucial to correctly predict the E200D SNP functionality in PykA and to anticipate its mild impact under the tested conditions.

While non-metabolic SNPs are dominant in MTBC strains, our analysis shed light on the underlying genetic basis for metabolic diversity and enabled a systematic characterization of divergent evolution in MTBC lineages. In addition, we showed that the approach could open the door to strain-specific drug treatments. As a proof of principle, we demonstrated significant differences in the susceptibility to PAS among MTBC strains. These experiments were motivated by our model-based results, and the strains’ differential sensitivities could be rationally linked to predicted functional SNPs in enzymes involved in the utilization and recycling of folate cofactors. Such predictions can guide the genome-wide analysis of drug-resistant MTBC, possibly identifying natural genetic variants that facilitate emergence of resistance. More generally, we envisage our model-based approach to enable a more systematic understanding of the key metabolic dependencies in pathogenic bacteria and to aid the development of selective treatments that exploit metabolic vulnerabilities during infection.

## Materials and Methods

### Cultivation and sampling

Three biological replicates of the 18 employed MTBC strains^33^ were cultured in 50 ml conical tubes containing 15 ml of medium incubated at 37°C and shaken continuously on an orbital rotator. We used a modified 7H9 medium (BD) supplemented with 0.5% (w/v) pyruvate, 0.05% (v/v) tyloxapol, 0.2% (w/v) glucose, 0.5% (w/v) bovine serum albumin (Fraction V) and 14.5 mM NaCl. Compared to the usual composition of 7H9 we omitted glycerol, tween 80, oleic acid and catalase from the medium. We monitored growth by determining optical density measured at 600 nm (OD_600_). To test pykA SNP functionality, we inoculated 10 ml of appropriate medium (with or without pyruvate) with ~10^7^ bacterial cells of N0004, N0091, N1176, N1274 or N1283 (starting OD_600_ < 0.01). We incubated the cultures on a rotatory shaker at 37°C and quantified their growth by optical density after 10 days. For analysis of the exometabolome, we periodically withdrew 1 ml aliquots, pelleted the cells by centrifugation (10,000 × *g*, 5 min), filtered the supernatant through 0.22 µm syringe filters twice to remove viable bacteria and kept the resulting supernatant at −80°C until mass spectrometric analysis. For the analysis of the intracellular metabolome, we harvested 3 OD_600_ unit equivalents of cells during mid-exponential growth phase (OD_600_ = 0.50 ± 0.10) using rapid filtration through a 0.45 μm filter. We washed the pellets with 2 ml of 75 mM ammonium carbonate buffer (pH 6.6, pre-warmed to 37°C). Metabolites were extracted by transferring the filter with cell pellets into a 15 ml conical tube containing 7.5 ml of ice-cold acetonitrile:methanol:water (2:2:1) solution, and incubated at −20°C for 1-4 hours. After incubation we filtered the extraction solution through 0.22 µm syringe filters twice to remove viable bacteria and kept the resulting supernatant was at −80°C until mass spectrometric analysis.

### Genome sequencing

Details on MTBC genome sequencing and bioinformatics analysis for sequence read alignment and variant determination are described in detail here^36^.

### Metabolomics profiling

Intracellular extracts and supernatant aliquots were directly injected into an Agilent 6550 time-of-flight mass spectrometer (ESI-iFunnel Q-TOF, Agilent Technologies)^35^. The platform consists of an Agilent Series 1100 LC pump coupled to a Gerstel MPS2 autosampler and an Agilent 6550 Series Quadrupole Time of Flight mass spectrometer (Agilent, Santa Clara, CA). Mass spectra were recorded from m/z 50 to 1000 using the highest resolving power (4 GHz HiRes). Detected ions were matched to a list of metabolites based on the corresponding molar mass. We compiled a comprehensive list of metabolites from the following genome-scale metabolic models: an automatically constructed *M. tuberculosis* H37Rv model from the Model SEED resource (Seed83332.1)^56^, a manually curated *M. tuberculosis* H37Rv model (sMtb)^57^, an *E. coli* K12 model (iJO1366)^58^, and a *Mycobacterium smegmatis* mc^2^155 model from the BioModels database (BMID000000141548, https://www.ebi.ac.uk/biomodels-main)^59^. The chemical formula of each metabolite was used to calculate the deprotonated monoisotopic molecular mass. Detected ions within a mass tolerance difference of less than 0.003 Da were associated to the nearest reference metabolites. This method is not able to separate compounds with similar m/z and relies on direct ionization without LC separation. Spectral data processing allowed annotation of 294 intracellular and 272 extracellular metabolites.

### Growth, uptake, and secretion rates

Growth rates and their 95% confidence intervals were estimated by fitting a line to log OD_600_ as a function of time, pooling data from all available biological replicates for each strain (three or four for MTBC strains, one for *P. aeruginosa*). Doubling times, *t*_d_, were calculated from the growth rates, *μ*:

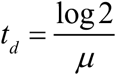

Growth rates for five MTBC strains with and without supplemented pyruvate were calculated in the same way (one replicate for each strain and condition).

Relative metabolite uptake and secretion rates were inferred from dynamic exometabolomes. For each annotated metabolite and each strain, linear models relating ion intensity to growth-normalized time (OD_600_ divided by growth rate) were fitted separately to biological replicates. If the slope of at least one linear model was significantly different from zero (*t*-test, *p* ≤ 0.05), the strain was considered capable of taking up (negative slope) or secreting (positive slope) the metabolite. Notably, we do not discriminate for the possibility that changes in the concentration of extracellular metabolites are mediated by secreted proteins. For MTBC strains, we also obtained bounds on relative uptake and secretion rates for each metabolite from ratios of significant slopes between different strains. We used the mean relative rate of each annotated metabolite in each strain to compute *L*^2^ (Euclidean) distances between the exometabolomes of pairs of strains, normalizing the uptake and secretion rates of each metabolite by their *L*^2^ norm.

### Strain-specific metabolic models

A published genome-scale metabolic model of *M. tuberculosis* H37Rv (iNJ661)^37^ was used as a template and customized for each strain. Glycerol was removed from the biomass composition and 95% confidence intervals of experimental growth rates were used to set upper and lower bounds on the flux of the biomass reaction. Bounds were set on boundary reactions to allow uptake of O_2_ and all defined components of the medium (biotin, Ca^2+^, citrate, Cl^-^, Cu^2+^, Fe^2+^, Fe^3+^, glucose, glutamate, H^+^, H_2_O, Mg^2+^, Na^+^, NH_4_^+^, PO_4_^3-^, pyruvate, pyridoxin, and SO_4_^2-^). We allowed secretion of all metabolites with a significant positive slope in the exometabolome data and of the simple inorganic compounds CO, CO_2_, H^+^, H_2_, H_2_O, H_2_S, and NH_4_^+^. To consider limited carbon sources in the growth medium (pyruvate, glucose, glutamate, and citrate), we set theoretical upper bounds on uptake rates:

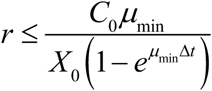

where *r* is the uptake rate, *C_0_* is the initial metabolite concentration, *µ*_min_ is the lower bound of the experimental growth rate, *X*_0_ is the estimated initial cell concentration, and Δ*t* is the duration of the experiment. Genes with sequencing coverage lower than five reads or 10% of the average coverage of the genome were considered to be deletions and removed from the model (**Supplementary Table 7**) along with all reactions requiring enzymes encoded by the deleted genes. We also removed blocked reactions that could never carry flux under the given constraints along with all metabolites participating only in blocked reactions.

### Statistical analysis of metabolome data

A One-Way Analysis of Variance was used to find metabolites exhibiting significant secretion/uptake difference across lineages. 18 strains and 6 lineages were analysed, yielding a between-groups degrees of freedom of 12. Estimated p-values were corrected for multiple tests using the procedure described in^60^.

### Structural sensitivity analysis

To predict the direct and indirect (network-mediated) effects of SNPs on metabolic fluxes, we used structural sensitivity analysis^39^ to determine the strain-specific sensitivity matrices **S**_*i*_. Originally, these matrices predict the response of a flux distribution to a perturbed flux through a target reaction, assuming minimal adjustments of fluxes. Here, we extended the concept to account for gene-level perturbations that may affect multiple reactions. We pre-processed each strain-specific model by removing the biomass reaction, by making all reactions reversible, and by aligning all metabolic reactions associated with each gene in the model (via reversing reaction directions such that the overlap of metabolites on each side of the reaction equations was maximal). We assumed that perturbation of a gene implied identical flux perturbations for all reactions associated with this gene. If such an identical flux perturbation was not feasible (e.g., because flux bounds were violated), we found the closest feasible perturbation by minimizing the *L*^2^ distance to the identical flux perturbation. Next, we fixed the perturbed fluxes and computed the minimal network response by minimizing the *L*^2^ norm of all fluxes not directly linked to the perturbed gene. If the minimal response was a thermodynamically infeasible loop^61^, we added constraints and variables to specifically disable this loop^62^ and re-computed a minimal response until the minimal loopless response was found. Finally, we added the gene-level perturbations and their minimal loopless responses (normalized by the mean of non-zero fluxes in each perturbation) as columns to the strain-specific sensitivity matrix, **S**_*i*_.

### Functional SNP prediction

The strain-specific models were combined into a single constraint-based model and the following balance constraints were added for each strain *i*:

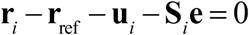

These balances couple strains to each other and ensure that all differences between fluxes in strain *i*, **r**_*i*_, and fluxes in a reference strain, **r**_ref_, are accounted for by unexplained effects, **u**_*i*_, and the effects of non-synonymous SNPs on enzymatic activities, **e**. The strain-specific sensitivity matrix **S**_*i*_ transforms SNP effects into fluxes. We added bounds on relative uptake and secretion rates from exometabolomes as well as bounds preventing the flux effects of SNPs from being larger than the affected reference fluxes. We solved the model by sequentially minimizing the *L*^1^ norm of (i) unexplained flux effects, (ii) reference fluxes, and (iii) SNP effects. We used the minimum obtained from each step to constrain the model before proceeding to the next step.

Flux variability analysis (FVA)^40^ was used to classify SNPs as functional or non-functional. Each SNP effect was minimized and maximized while requiring unexplained flux effects to be at the minimum obtained from the first minimization step. A SNP was classified as functional if its minimal effect was larger than a chosen flux threshold (*α* = 10^-4^ mmol gDW^-1^ h^-1^) and non-functional otherwise. The relative flux effect of a functional SNP was determined by dividing its minimal effect by the mean flux of reactions catalyzed by the affected enzyme in the reference flux distribution obtained after the third minimization step.

We obtained SIFT scores for SNPs relative to the laboratory strain H37Rv from SIFT4G^28^ (http://sift.bii.a-star.edu.sg/sift4g/) and filtered out predictions with median sequence information above 3.25 to avoid prediction based on closely related sequences. SNPs with SIFT score (probability of amino acid change being tolerated) below the recommended threshold of 0.05 were classified as functional according to SIFT.

### Intracellular metabolome analysis

For each non-synonymous SNP and each metabolite in the intracellular metabolome participating in a reaction catalyzed by the affected enzyme, the mean intracellular level in strains with the SNP was divided by the mean intracellular level in strains without the SNP. The resulting relative intracellular levels were used to compute the mean relative intracellular levels of functional and non-functional SNPs (as classified by our model and SIFT), and bootstrapping with 1,000 samples was used to compute 95% confidence intervals. Random distributions of mean relative intracellular intensities were obtained by randomizing the strain-assignment of SNPs 1,000 times while preserving the number of strains affected by each SNP.

### Synthetic lethality prediction

For each strain and each SNP classified as functional in that strain, the gene containing the SNP was deleted in the strain-specific model. If the gene was not essential, all other non-essential genes were deleted one by one. Pairwise deletions that prevent biomass production were identified as synthetic lethal interactions.

### Antibiotic sensitivity analysis

The sensitivity of strains to PAS and OFX was determined as described elsewhere^63^. Briefly, we prepared 96-well plates such that they contained a 2-fold serial dilution of a drug of interest (e.g., PAS, starting from a concentration of 4 μg/ml) in 90 μl of our modified 7H9 medium. We added 10 μl of bacterial culture adjusted to a starting OD_600_ of 0.02. Plates were incubated for 10 days at 37°C after which time we added 10 μl of 0.02% alamar blue solution and incubated for a further 24 h. After 24 h we added 100 μl of 3.7% formalin solution to kill the bacteria and scored the plates by measuring the fluorescence (*λ*_ex_ = 545 nm, *λ*_em_= 590 nm) using a Molecular Devices spectraMAX microplate reader.

For each drug and each strain, fluorescence signals were normalized by (1) subtracting the signal of the drug without any cells and (2) dividing by the signal of cells without drug. For each biological replicate, the maximum drug effect was quantified as one minus the ratio of fluorescences at the highest and lowest drug dose, respectively. The mean maximum effect was calculated for each lineage with 95% confidence intervals obtained from bootstrapping with 1,000 samples.

### Software

Models were built in Python using COBRApy^64^ and optimization problems were solved with the Gurobi Optimizer 7.5.2 (Gurobi Optimization, LLC, Beaverton, OR, USA). Inference of metabolite uptake and secretion and all steps of mass spectrometry data processing and analysis were performed in MATLAB R2018 (The MathWorks, Inc., Natick, MA, USA). Remaining data analysis was performed in Python.

### Data and code availability

All data presented in this study are available as supplementary materials. The model and the code needed to solve it and reproduce our results are available at https://gitlab.com/csb.ethz/MtbSNP.

## Acknowledgments

We thank Dr. Thomas Liphardt, Dr. Hans-Michael Kaltenbach, and Charlotte Ramon for helpful contributions to the analysis. This work was supported by the Swiss National Science Foundation (grants 310030_166687, IZRJZ3_164171, IZLSZ3_170834, and CRSII5_177163), the European Research Council (309540-EVODRTB), and SystemsX.ch (TbX).

## Contributions

M.Za., O.Ø. and J.S. designed the computational analysis. S.G., U.S. S.B and A.T. designed the experiments. S.B., A.T and J.F. performed the experiments. M.Za. and M.Zi. performed the metabolomics measurements. O.Ø. and M.Za. analyzed the data. M.Za, O.Ø., A.T, U.S., and J.S. wrote the manuscript. All authors contributed to preparing the manuscript.

## Competing interests

The authors declare that they have no competing interests.

## Supplementary Materials

Supplementary Figs. 1-6, Supplementary Tables 1-7.

