## Supplementary Materials for "Model-based integration of genomics and metabolomics reveals SNP functionality in *Mycobacterium tuberculosis*"

### Contents

**1 Supplementary figures** **2**

**2 Detailed model description** **6**

### 1 Supplementary figures

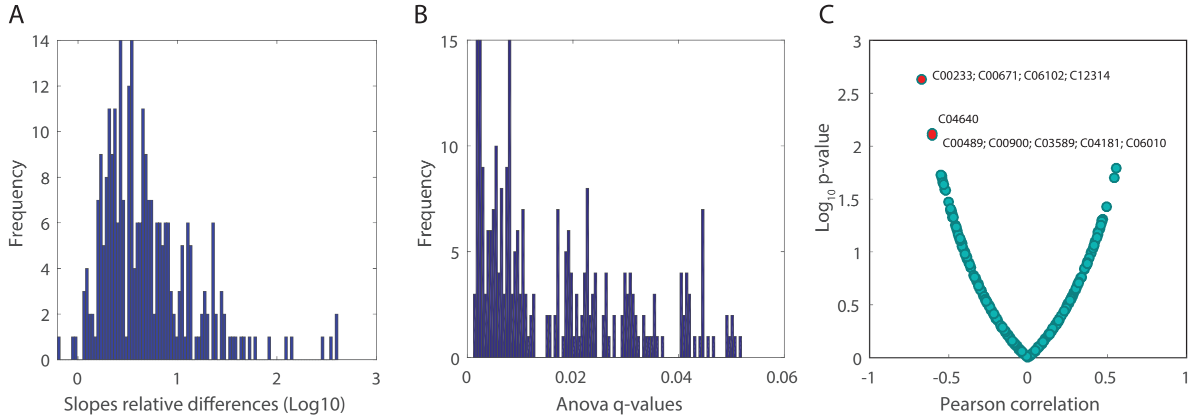


**Figure 1:** Statistical analysis of differences in exometabolome data. (A) Histogram of largest differences between uptake/secretion rates of metabolites across strains. For each metabolite the maximum fold change between all possible strain pairs is reported. (B) Distribution of corrected ANOVA $p$-values ($q$-values). (C) In this scatter plot each dot represents an annotated ion and its correlation/significance between relative metabolite levels and growth rates across MTBC strains.

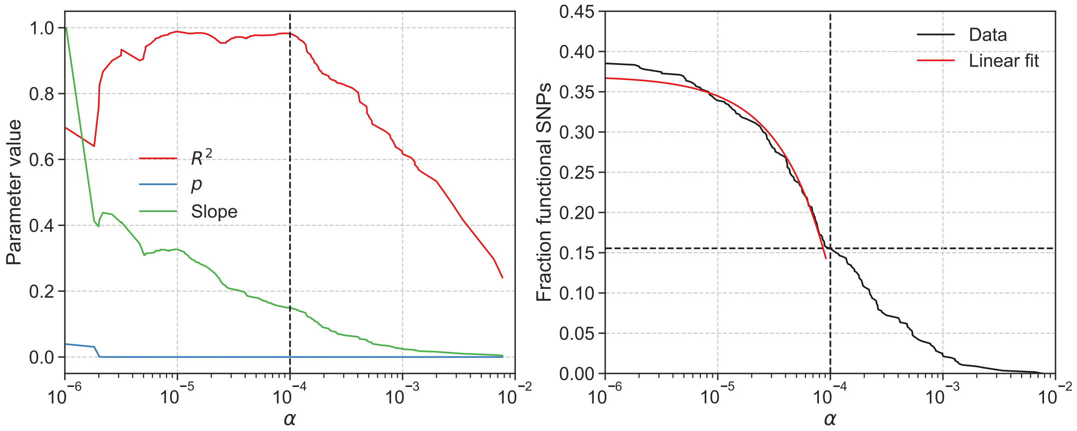


**Figure 2:** We determined the threshold for SNP classification, $\alpha$, by fitting linear models to the fraction of functional SNPs as a function of $\alpha$, starting from the smallest $\alpha$ and increasing it through its possible values, one by one. The coefficient of determination, $R^2$, the probability of non-zero slope $p$, and the relative slope are shown on the left. The chosen $\alpha = 10^{-4}$ (indicated by vertical dashed line) is very close to the last peak of $R^2$ before it starts monotonously decreasing with increasing $\alpha$. On the right, the fraction of funtional SNPs is shown as a function of $\alpha$ along with the linear fit for $\alpha = 10^{-4}$. The fraction of functional SNPs for $\alpha = 10^{-4}$ is indicated by a dashed horizontal line.

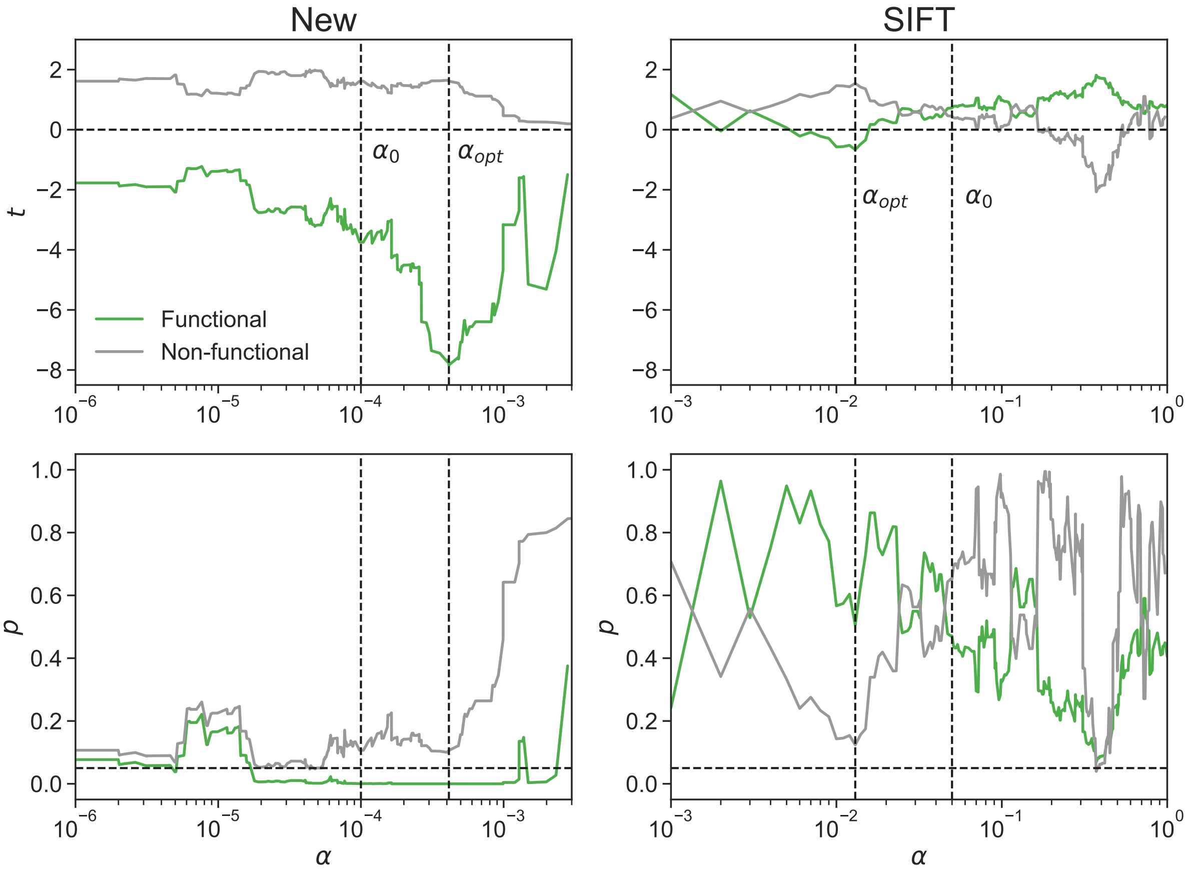


**Figure 3:** Results of $t$-tests comparing the mean relative intracellular level of directly affected metabolites to zero for functional and non-functional SNPs classified by our new method and SIFT. The $t$-statistic and $p$-value are shown as a function of the classification threshold, $\alpha$. Vertical dashed lines indicate the threshold used for classification ($\alpha_0$) and a stricter threshold that optimizes separation of functional and non-functional SNPs ($\alpha_{\text{opt}}$). Horizontal dashed lines indicate $t = 0$ and $p = 0.05$.

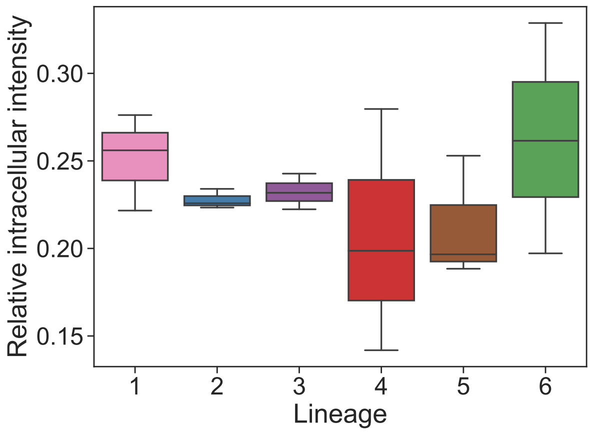


**Figure 4:** Relative intracellular levels of methylglyoxal by lineage.

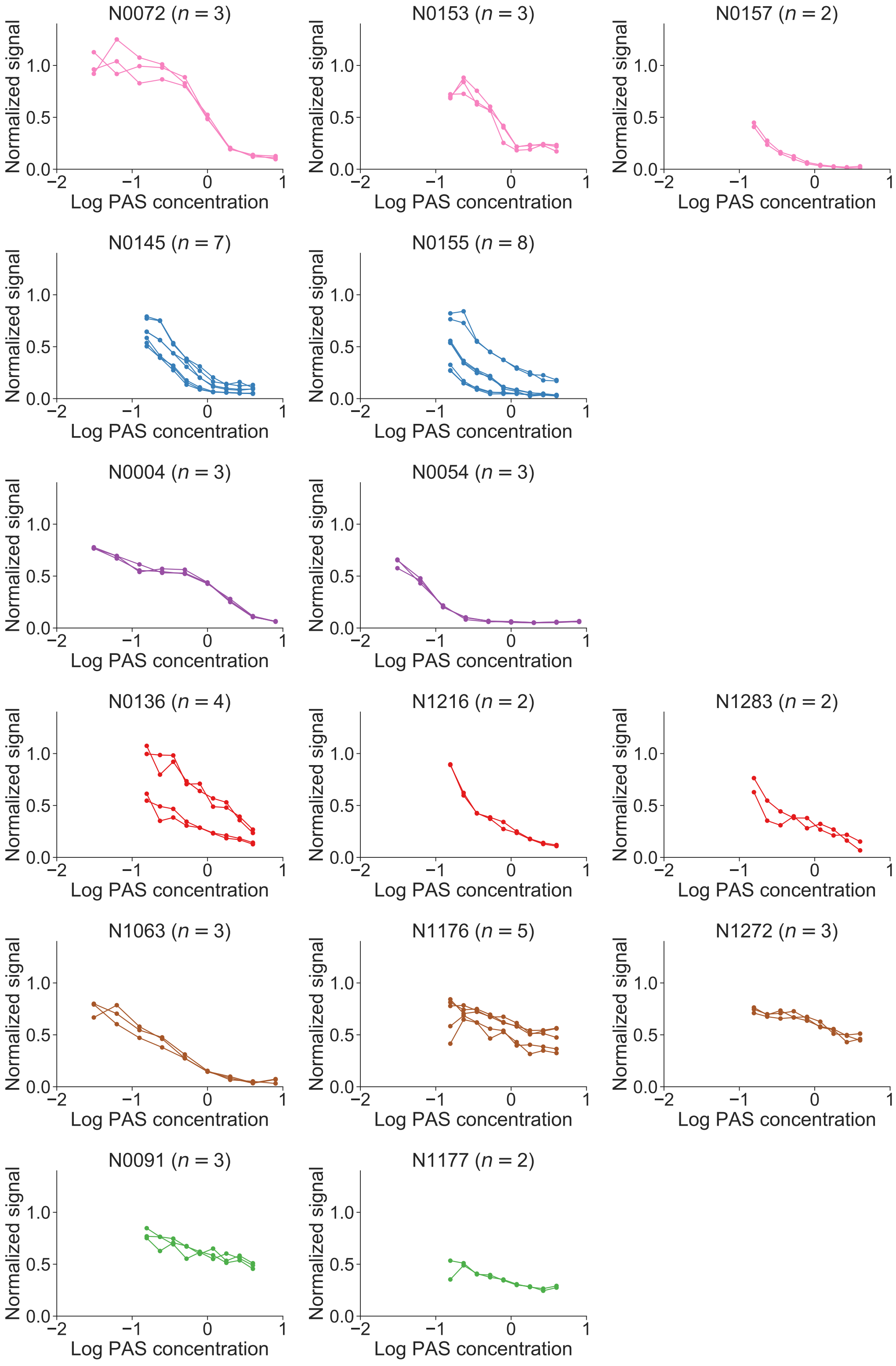


**Figure 5:** Fluorescence signal reflecting cell concentrations as a function of log *para*-aminosalicylic acid (PAS) concentration. Strains are ordered by lineage from top to bottom. Number of replicates are indicated for each strain.

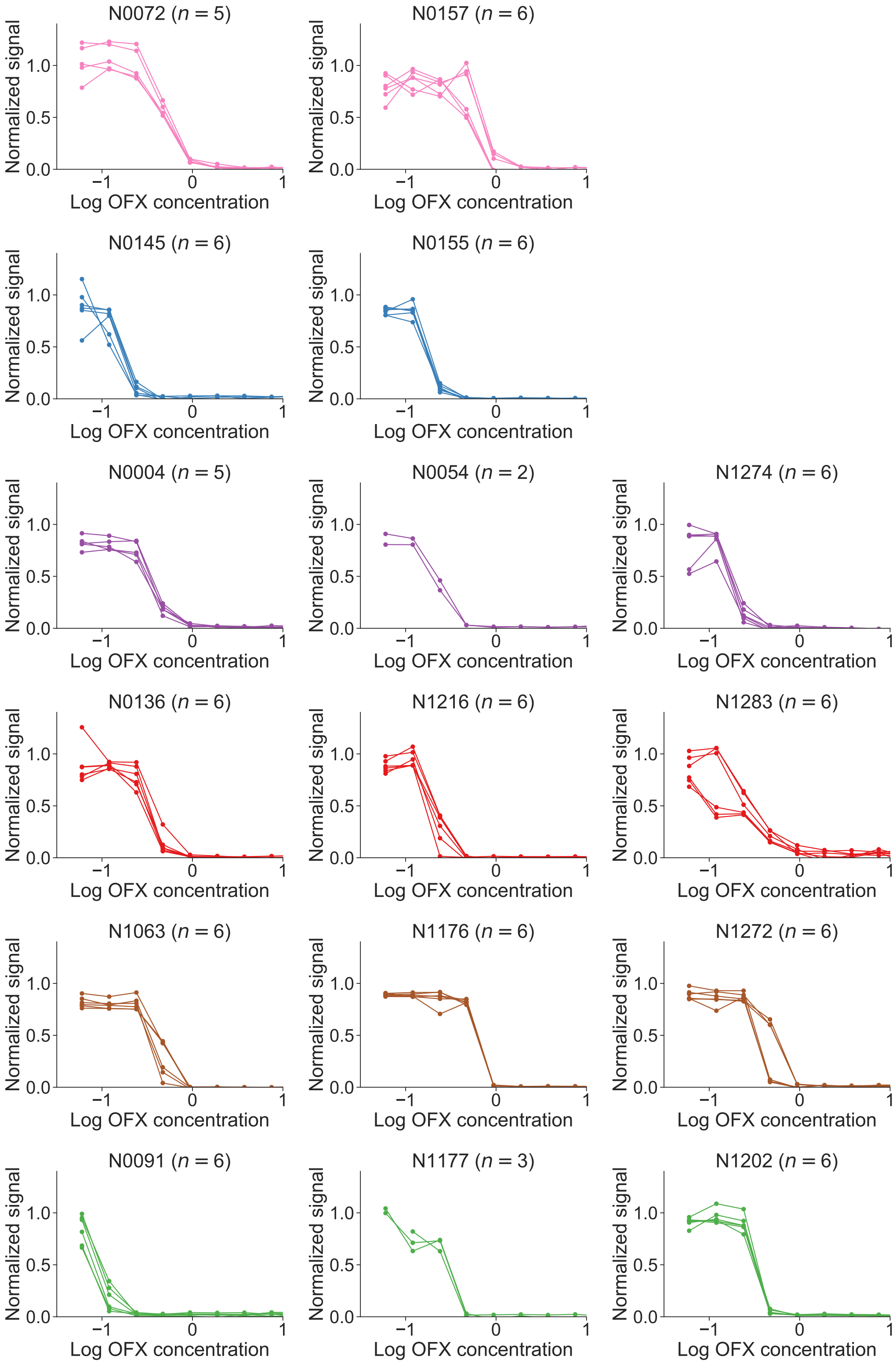


**Figure 6:** Fluorescence signal reflecting cell concentrations as a function of log ofloxacin (OFX) concentration. Strains are ordered by lineage from top to bottom. Number of replicates are indicated for each strain.

### 2 Detailed model description

The variables in the model are the fluxes in each strain $i$, $\mathbf{r}_i$, fluxes in a reference strain, $\mathbf{r}_{\text{ref}}$, unexplained flux effects, $\mathbf{u}_i$, and SNP flux effects, $\mathbf{e}$.

We represent the metabolic network of strain $i$ by its stoichiometric matrix, $\mathbf{N}_i$, and make the quasi-steady-state assumption:

$$\mathbf{N}_i \mathbf{r}_i = 0 \tag{1}$$

Any flux difference between strains (any deviation from the reference fluxes) must be accounted for by unexplained effects and SNP effects:

$$\mathbf{r}_i - \mathbf{r}_{\text{ref}} - \mathbf{u}_i - \mathbf{S}_i \mathbf{e} = 0 \tag{2}$$

SNP effects are translated into fluxes by the strain-specific SNP sensitivity matrix, $\mathbf{S}_i$, in which the columns are SNPs, the rows are fluxes, and each element is the effect of a normalized SNP perturbation on a flux in strain $i$.

Fluxes in all strains $i$ as well as the reference strain have upper and lower bounds:

$$\mathbf{r}_i^{\min} \leq \mathbf{r}_i \leq \mathbf{r}_i^{\max} \tag{3}$$

We also constrain the boundary flux ratio for metabolite $m$ between pairs of strains, $i$ and $j$:

$$\alpha_{i,j}^m \leq \frac{\sum_{k \in \mathcal{B}_i^m} b_i^k}{\sum_{k \in \mathcal{B}_j^m} b_j^k} \leq \beta_{i,j}^m \tag{4}$$

Here, $\mathcal{B}_i^m$ is the set of boundary fluxes in strain $i$ mapped to metabolite $m$ and $b_i^k$ is boundary flux $k$ in strain $i$. The ratio bounds, $\alpha_{i,j}^m$ and $\beta_{i,j}^m$, are inferred from dynamic exometabolomes.

Unexplained effects are unconstrained but SNP effects can only be negative because we only allow deleterious SNPs:

$$\mathbf{e} \leq 0 \tag{5}$$

We predict SNP effects by solving three sequential linear programming problems. First, we minimize the $L^1$ norm of unexplained effects:

$$\begin{array}{lll}
\text{minimize} & \|\mathbf{u}\|_1 & \\
\text{subject to} & \mathbf{N}_i \mathbf{r}_i = 0 & \forall i \in \mathcal{S} \\
& \mathbf{r}_i - \mathbf{r}_{\text{ref}} - \mathbf{u}_i - \mathbf{S}_i \mathbf{e} = 0 & \forall i \in \mathcal{S} \\
& \alpha_{i,j}^m \leq \frac{\sum_{k \in \mathcal{B}_i^m} b_i^k}{\sum_{k \in \mathcal{B}_j^m} b_j^k} \leq \beta_{i,j}^m & \forall i \in \mathcal{S},\ \forall j \in \mathcal{S},\ \forall m \in \mathcal{M} \\
& \mathbf{r}_i^{\min} \leq \mathbf{r}_i \leq \mathbf{r}_i^{\max} & \forall i \in \mathcal{S} \\
& \mathbf{r}_{\text{ref}}^{\min} \leq \mathbf{r}_{\text{ref}} \leq \mathbf{r}_{\text{ref}}^{\max} & \\
& \mathbf{e} \leq 0 &
\end{array}$$

Here, $\mathcal{S}$ is the set of strains and $\mathcal{M}$ is the set of metabolites in the exometabolome.

Second, we fix the $L^1$ norm of unexplained effects to the minimum obtained from the previous step and minimize the $L^1$ norm of reference fluxes:

$$\begin{array}{lll}
\text{minimize} & \|\mathbf{r}_{\text{ref}}\|_1 & \\
\text{subject to} & \mathbf{N}_i\mathbf{r}_i = 0 & \forall i \in \mathcal{S} \\
& \mathbf{r}_i - \mathbf{r}_{\text{ref}} - \mathbf{u}_i - \mathbf{S}_i\mathbf{e} = 0 & \forall i \in \mathcal{S} \\
& \alpha_{i,j}^m \leq \frac{\sum_{k\in\mathcal{B}_i^m} b_i^k}{\sum_{k\in\mathcal{B}_j^m} b_j^k} \leq \beta_{i,j}^m & \forall i \in \mathcal{S},\ \forall j \in \mathcal{S},\ \forall m \in \mathcal{M} \\
& \mathbf{r}_i^{\min} \leq \mathbf{r}_i \leq \mathbf{r}_i^{\max} & \forall i \in \mathcal{S} \\
& \mathbf{r}_{\text{ref}}^{\min} \leq \mathbf{r}_{\text{ref}} \leq \mathbf{r}_{\text{ref}}^{\max} & \\
& \mathbf{e} \leq 0 & \\
& \|\mathbf{u}\|_1 = \|\mathbf{u}\|_1^{\min} &
\end{array}$$

Finally, we fix the $L^1$ norms of unexplained effects and reference fluxes to the minima obtained from the two previous steps and minimize the $L^1$ norm of SNP effects:

$$\begin{array}{lll}
\text{minimize} & \|\mathbf{e}\|_1 & \\
\text{subject to} & \mathbf{N}_i\mathbf{r}_i = 0 & \forall i \in \mathcal{S} \\
& \mathbf{r}_i - \mathbf{r}_{\text{ref}} - \mathbf{u}_i - \mathbf{S}_i\mathbf{e} = 0 & \forall i \in \mathcal{S} \\
& \alpha_{i,j}^m \leq \frac{\sum_{k\in\mathcal{B}_i^m} b_i^k}{\sum_{k\in\mathcal{B}_j^m} b_j^k} \leq \beta_{i,j}^m & \forall i \in \mathcal{S},\ \forall j \in \mathcal{S},\ \forall m \in \mathcal{M} \\
& \mathbf{r}_i^{\min} \leq \mathbf{r}_i \leq \mathbf{r}_i^{\max} & \forall i \in \mathcal{S} \\
& \mathbf{r}_{\text{ref}}^{\min} \leq \mathbf{r}_{\text{ref}} \leq \mathbf{r}_{\text{ref}}^{\max} & \\
& \mathbf{e} \leq 0 & \\
& \|\mathbf{u}\|_1 = \|\mathbf{u}\|_1^{\min} & \\
& \|\mathbf{r}_{\text{ref}}\|_1 = \|\mathbf{r}_{\text{ref}}\|_1^{\min} &
\end{array}$$
